# Asymmetric induction of IL-23R by CpG and IL-15 in proliferative CLL fractions highlights intraclonal heterogeneity in chronic lymphocytic leukemia

**DOI:** 10.1101/2025.08.27.672649

**Authors:** Martina Cardillo, Nadia Bertola, Fabiana Ferrero, Rosanna Massara, Maria Cristina Capra, Daniele Reverberi, Monica Colombo, Antonino Neri, Fortunato Morabito, Franco Fais, Giovanna Cutrona

**Author notes:** Authors contributed equally to the work (Franco Fais, Giovanna Cutrona).

## Abstract

Chronic lymphocytic leukemia (CLL) is sustained by complex interactions with the microenvironment, which provides signals for leukemic cell survival and proliferation. Among these, cytokines of the IL-12 family have emerged as relevant regulators of immune responses and tumor biology, yet their contribution to CLL remains incompletely defined. Previous studies showed that CLL cells can acquire responsiveness to IL-23 after T cell–dependent stimulation with CD40L, raising the question of whether T cell–independent signals can similarly induce functional receptor expression.

Here we investigated the effect of CpG oligodeoxynucleotides, alone or in combination with interleukin-15 (IL-15), on IL-12 family receptor expression in CLL cells. We found that CpG + IL-15 stimulation significantly increased IL-23R and IL-12Rβ1 expression, while IL-12Rβ2 remained largely unresponsive. As a consequence, the complete IL-23 receptor complex was robustly induced, whereas IL-12 receptor assembly was only marginally enhanced. This skewing toward IL-23 rather than IL-12 signaling suggests that innate immune stimuli preferentially promote pathways supporting inflammation and survival, while limiting tumor-suppressive IL-12 responsiveness.

Analysis of intraclonal heterogeneity revealed an asymmetric distribution of IL-23R among CXCR4/CD5-defined subfractions: the proliferative fraction, representing recently divided cells, expressed higher levels of IL-23R compared to the resting fraction. These findings suggest that IL-23 responsiveness is particularly enriched in the proliferating compartment of the leukemic clone.

Overall, our results indicate that CpG and IL-15 stimulation drive a selective expansion of IL-23 signaling capacity in CLL, with preferential engagement of proliferative subfractions. This imbalance between IL-23 activation and insufficient IL-12Rβ2 induction may represent a critical pathogenic mechanism and a potential therapeutic target.

## INTRODUCTION

Chronic lymphocytic leukemia (CLL) is the most common leukemia in adults in Western countries and is characterized by the clonal accumulation of mature CD5^+^ B lymphocytes in the peripheral blood, bone marrow, and secondary lymphoid organs (Hallek et al., 2018). Early reviews emphasized that CLL pathogenesis is strongly shaped by chronic antigenic stimulation and by the biology of the B-cell receptor (BCR), which plays a central role in leukemic cell survival and clonal evolution (Annu Rev Immunol, 2003). Subsequent analyses also redefined CLL as a dynamic disease in which leukemic cell turnover and continuous trafficking between lymphoid tissues and peripheral blood, rather than passive accumulation alone, underlie disease progression (Hematology ASH Educ Program, 2006). Although many patients present with an indolent disease course and may not require therapy for years, a significant subset exhibits aggressive progression necessitating early treatment. Disease heterogeneity reflects both intrinsic features of the leukemic clone and extrinsic influences from the microenvironment, which provides survival, activation, and proliferative signals necessary for disease evolution (Burger et al., 2009; Ghia et al., 2008; Herishanu et al., 2011; Kipps et al., 2017).

In addition to interpatient variability, CLL displays intraclonal complexity, whereby the leukemic population comprises subfractions with distinct proliferative histories and biological properties (Calissano et al., 2011). A well-established approach to dissect this complexity relies on the differential surface expression of CD5 and the chemokine receptor CXCR4, which allows the identification of fractions along a proliferation/quiescence continuum: CXCR4^dim/CD5^bright cells represent recently divided, proliferation-competent elements that have recently exited lymphoid tissues, whereas CXCR4^bright/CD5^dim cells correspond to older, quiescent cells that have resided longer in the peripheral blood. Intermediate fractions reflect transitional stages in this trafficking cycle. These subpopulations differ in telomere length, gene expression programs, activation marker profile, and responsiveness to microenvironmental cues. Importantly, the CXCR4^dim/CD5^bright fraction is enriched for cells recently stimulated in proliferation centers within lymph nodes, suggesting that they may be susceptible to further activating signals upon re-entry into supportive niches.

Among the mediators shaping CLL biology, the interleukin-12 (IL-12) cytokine family plays a central role in immune regulation and inflammation. This family includes IL-12, IL-23, IL-27, and IL-35, all heterodimeric cytokines sharing subunits and receptor components (Trinchieri, 2003). IL-12 signals through a heterodimeric receptor composed of IL-12Rβ1 and IL-12Rβ2 chains, while IL-23 signals via a receptor consisting of IL-12Rβ1 paired with the IL-23R chain (Oppmann et al., 2000). IL-12 typically promotes Th1 immune responses and enhances cytotoxicity (Szabo et al., 2003), whereas IL-23 is involved in bridging innate and adaptive immunity, supporting Th17 differentiation, and sustaining chronic inflammation (Langrish et al., 2005).

In normal B-cell physiology, expression of the IL-23 receptor complex has been detected in early B lymphocytes, germinal center B cells, and plasma cells (Kotenko et al., 2001; Parham et al., 2002), but not typically in mature circulating B cells. In malignant contexts, IL-23R expression has been described in acute lymphoblastic leukemia, follicular lymphoma, diffuse large B-cell lymphoma, and multiple myeloma (Tangye et al., 2001; Tangye et al., 2003; Lee et al., 2014).

Previous work by our group demonstrated that circulating CLL cells often exhibit an “incomplete” IL-23 receptor phenotype, with expression of IL-23R in the absence of IL-12Rβ1. Notably, CD40/CD40L interactions provided by activated T cells can induce CLL cells to express the complete IL-23 receptor complex, including IL-12Rβ1 (Cutrona et al., 2018). Under these T cell–dependent activation conditions, CLL cells not only become responsive to IL-23 but also produce IL-23 themselves, suggesting an autocrine/paracrine stimulatory loop (Cutrona et al., 2018). In previous analyses, the expression of IL-12 and IL-23 receptor chains was assessed primarily in unstimulated CLL cells, both in bulk populations and, in selected cases, within the distinct CXCR4/CD5-defined intraclonal fractions.

Given the role of CD40L in enabling IL-12Rβ1 expression, it remains unresolved whether T cell– independent activation can similarly promote assembly of a functional IL-12/IL-23 receptors complex in CLL cells. CpG oligodeoxynucleotides, acting as TLR9 agonists, are known to activate CLL cells directly and induce proliferation, survival, NF-κB signaling, and cytokine production (Jahrsdörfer et al., 2001; Longo et al., 2007; Decker et al., 2000; Chen et al., 2015). Moreover, combining CpG with cytokines such as interleukin-15 (IL-15) can further enhance activation signals (Mongini et al., 2015). In the present study, we investigated whether stimulation with CpG or CpG + IL-15 can induce the expression of certain IL-12 family receptor chains in CLL cells, including within the different CXCR4/CD5-defined intraclonal fractions. This approach enables us to determine whether activation potential is uniformly distributed across the leukemic clone or enriched within subfractions characterized by features of recently divided, proliferation-competent leukemic cells.

## METHODS

### CLL cell samples

The study was approved by the Institutional Review Boards of Northwell Health and was conducted according to the principles of the World Medical Association Declaration of Helsinki. CLL patients were diagnosed as recommended, and all subjects provided written informed consent at enrollment. Furthermore, part of the experiments was conducted using peripheral blood samples from newly diagnosed patients with Binet stage-A CLL enrolled in the O-CLL1 protocol (clinicaltrial.gov identifier NCT00917540) (Morabito F et al 2013).

### CLL cells isolation

CLL cells from each patient’s PB were isolated by negative selection using RosetteSep Human B Cell Enrichment Cocktail (Stemcell Technologies, Vancouver, BC). Whole PB was incubated with the mixture, then diluted with 2% FBS in PBS and centrifuged over RosetteSep DM-L Density Medium (Stemcell Technologies). CLL samples initially purified by this technique were tested for purity by the Center for CLL Research. Cells were then resuspended in freezing solution and cryopreserved in liquid nitrogen. Samples from CLL patients that contained at least 95% of leukaemic cells were considered eligible for the study.

### CLL cells in vitro culture

Cell cultures of CLL cells were performed by seeding thawed cells in an enriched medium used for normal B cell replication in long-term85 cultures with added insulin/transferrin/selenium supplement (CAT #17-838Z; Lonza). Notably, this medium contains the reducing agent, 2-ME (5 × 10−5 M). The latter replaces an important function of bone marrow stromal cells in converting cystine to cysteine, which is needed for CLL uptake and use in the glutathione synthesis needed for retained viability. Fresh medium was prepared for each experiment using stock additives. Cultures were routinely established in 96-well round-bottom plates at 4 × 10^5 cells per 200-µl volume with duplicates for each culture condition. Recombinant human IL-15 (PeproTech Inc.) and CpG DNA TLR9 ligand (ODN-2006; Invivogen) were added at final culture concentrations of 15 ng/ml and 0.2 µM (1.5 µg/ml) for 72h, respectively.

### Detection of Cytokines Receptors in CLL by Flow Cytometry

Live cells were identified using LIVE/DEAD Fixable Stains for flow cytometry (LIVE/DEAD™ Fixable Violet Dead Cell Stain Kit or Far Red Dead Cell Stain Kit, Life Technologies). For surface membrane immunofluorescence, cells (2 × 10^5) in FACS buffer (PBS + 10% bovine serum albumin + 1% sodium azide) were incubated with primary antibody for 20 min at 4°C, followed by fixation with 0.1% formaldehyde in PBS. The same procedure and number of cells were used for the isotype control.

To detect the different chains, the following mAbs were used: IL23R (cat#FAB140019-100, R&D Systems), IL12Rß1 (cat #565043, BD Horizon), IL12Rß2 (cat #FAB1959C, R&D Systems). Data was acquired with a BD LSR Fortessa flow cytometer using the HTS plate reader and analyzed by FlowJo 10.6.2 version.

### Detection of Cytokines Receptors in CLL fractions by Flow Cytometry

PBMCs were thawed and stained for the surface markers CXCR4 APC (cat #306510, BioLegend), CD5 PE-Cy7 (cat #300622, BioLegend) and CD19 Pacific Blue (cat #48-0199-42, ebioscience). For the staining, cells (2 × 10^5) in FACS buffer (PBS + 10% bovine serum albumin + 1% sodium azide) were incubated with primary antibody for 20 min at 4°C, followed by fixation with 0.1% formaldehyde in PBS.

This staining allowed us to study 3 different fractions: Proliferative fraction (PF) CXCX4dim/CD5 bright; Resting fraction (RF) CXCR4 bright/CD5dim; Intermediate fraction (IF) CXCR4 dim/CD5dim87. For each of these fractions, we checked the expression of the receptors described above.

## RESULTS

### CpG+IL-15 stimulation induces IL-23 and IL-12 receptor complexes in CLL cells

We first examined whether the IL-23 receptor (IL23R) complex and IL-12 receptor (IL12R) complex could be induced in CLL cells following TLR9 activation. In line with previous observations (Cutrona et al., 2018), unstimulated ex vivo CLL cells expressed only low levels of the IL23R complex. Similarly, and consistent with earlier reports (Cutrona et al., 2018), the IL12R complex was virtually absent, primarily due to the consistently poor expression of the IL12Rβ2 chain (data not shown). To activate leukemic B cells, purified CLL cells were cultured for 72 hours with CpG oligodeoxynucleotides (CpG) alone or in combination with IL-15, a cytokine known to synergize with CpG in promoting CLL clonal expansion (Mongini et al., 2015). Both stimulation conditions led to a significant upregulation of the IL23R and IL12Rβ1 chains on the cell surface. In contrast, IL12Rβ2 expression was detected in only a small subset of cells, restricting the formation of the complete IL12R complex (Fig. 1A). Receptor complexes were quantified as double-positive populations for the relevant chains—IL23R^+^IL12Rβ1^+^ for the IL23R complex and IL12Rβ1^+^IL12Rβ2^+^ for the IL12R complex. After 72 hours of stimulation, both complexes were detectable, but IL23R complex expression was consistently higher than IL12R complex expression (Fig. 1B). This preferential induction of the IL23R complex reflected the limited expression of the IL12Rβ2 chain.

**Fig. 1.**
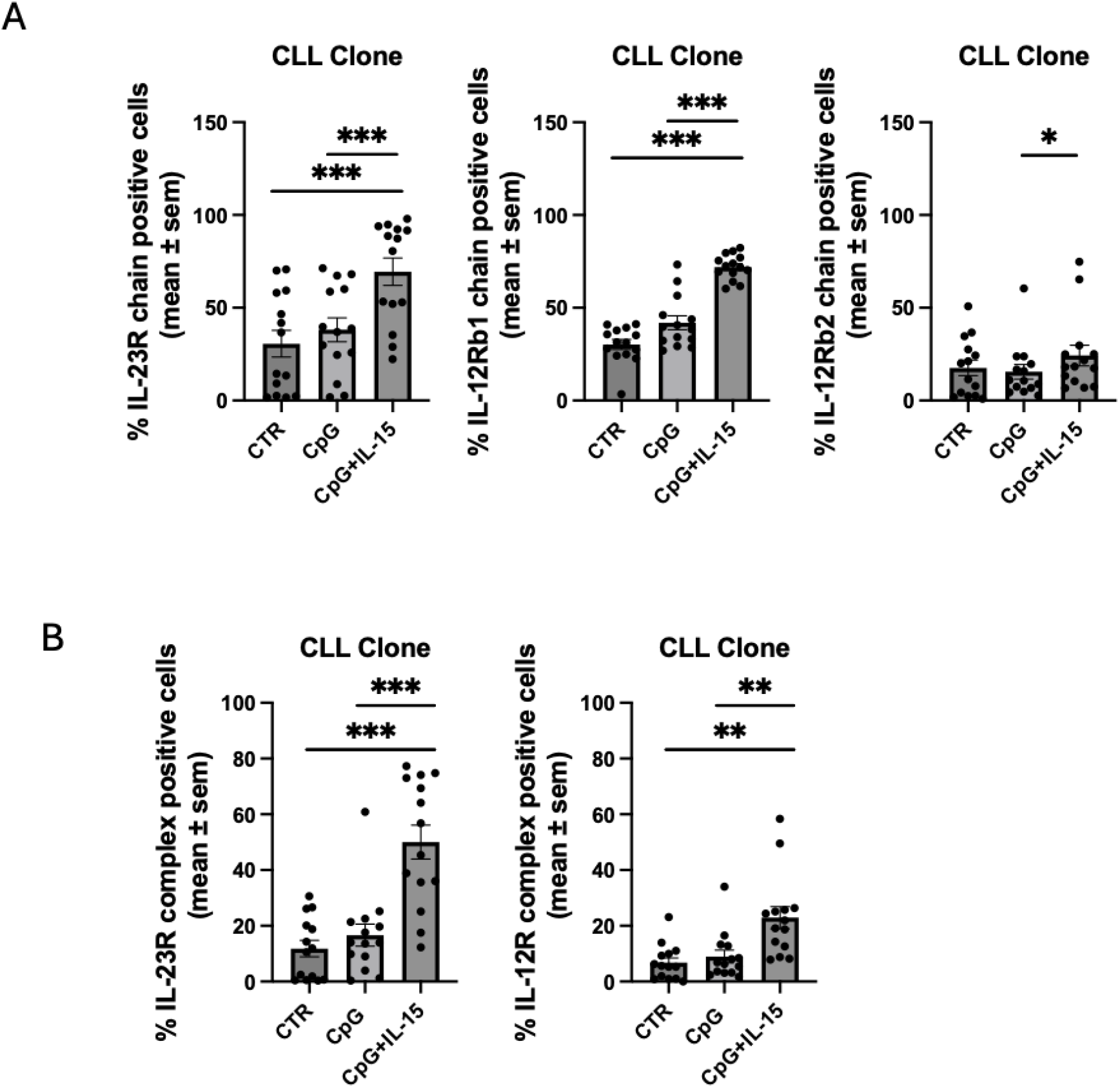
IL23R and IL12R expression in CLL. A. Expression of IL23R complex and IL12R complex on CLL cells after stimulation with CpG+IL-15 for 72h B. Expression of IL23R, IL12Rß1 and IL12Rß2 chains after 72h of stimulation in 14 CLL cell samples. The statistical significance of the difference is evaluated using the two-sided Wilcoxon signed-rank test. *p 0.05; **p 0.01; ***p 0.001; ****p 0.0001.

### CpG+IL-15 Stimulation Upregulates IL-23 and IL-12 Receptor Chains

To investigate whether IL-23 and IL-12 receptor expression varies with the activation or maturation stage of CLL cells, we analyzed their distribution across phenotypically distinct CLL fractions defined by Calissano et al. (2011). In this model of the CLL cell life cycle, leukemic cells residing in lymphoid tissue are retained in the stroma via CXCR4–CXCL12 interactions. Upon receiving activation signals, these cells enter the cell cycle, upregulate CD5, and internalize CXCR4. This CXCR4dim/CD5bright subset, termed the *Proliferating Fraction (PF)*, represents recently divided cells that have migrated from tissues into the blood, rather than cells actively dividing in circulation. Once in the peripheral blood, the lack of supportive microenvironmental signals leads to partial re-expression of CXCR4 and downregulation of CD5, giving rise to an *Intermediate Fraction (IF)* with a CXCR4int/CD5int phenotype. With additional time in circulation, CLL cells further increase CXCR4 and decrease CD5 expression, becoming part of the *Resting Fraction (RF)* (CXCR4bright/CD5dim). This phenotypic progression enables cells to re-enter lymphoid tissues by following the CXCR4–CXCL12 gradient. The gating strategy for these fractions is shown in Fig. 2A.

**Fig. 2.**
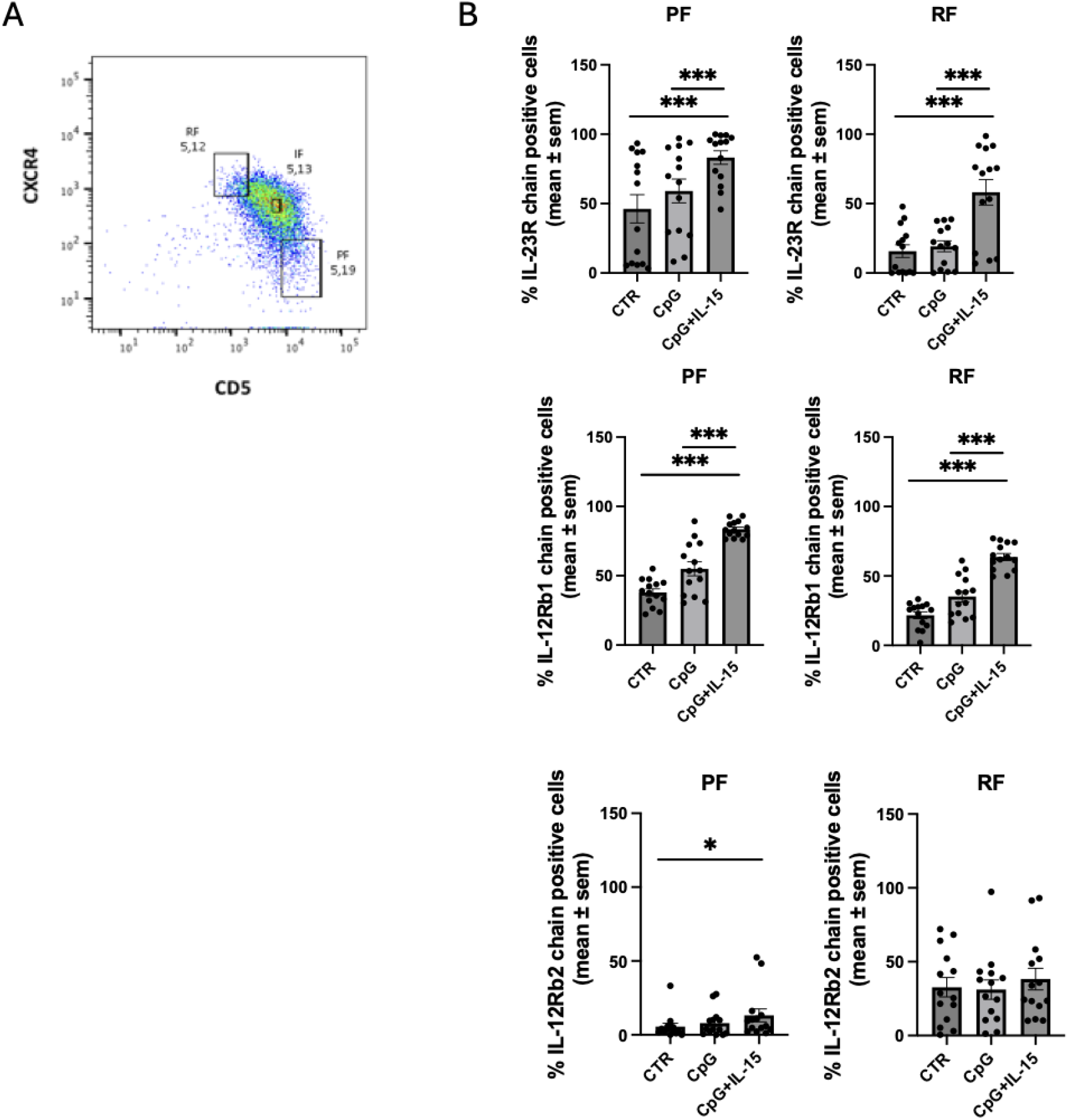
Expression of IL23R, IL12Rß1 and IL12Rß2 chains in CLL fractions. A. Example of gating strategy for CLL fractions B. Expression of IL23R, IL12Rß1 and IL12Rß2 chains after 72h of stimulation in PF and RF CLL fractions (n=14). The statistical significance of the difference is evaluated using the two-sided Wilcoxon signed-rank test. *p 0.05; **p 0.01; ***p 0.001; ****p 0.0001.

Using this framework, we assessed IL-23 and IL-12 receptor chain expression in PF and RF cells after 72 h of CpG+IL-15 stimulation. IL12Rβ1 expression increased in both PF and RF, but was consistently higher in PF. IL12Rβ2 remained low across all fractions, with a surprisingly slight increase in RF. In contrast, IL23R expression also rose after stimulation in both PF and RF, again with higher levels in PF compared to RF. These results indicate that the more recently divided PF cells are more responsive to CpG+IL-15, showing stronger induction of both IL12Rβ1 and IL23R, whereas IL12Rβ2 remains scarcely induced (Fig. 2B).

When we examined the expression of the complete receptor complexes (IL-12R and IL-23R) in PF and RF under each condition, we found that IL-12R levels did not differ significantly between fractions in CTR, CpG, or CpG+IL-15 cultures. By contrast, IL-23R expression was significantly higher in PF compared to RF after both CpG and CpG+IL-15 stimulation, further supporting a preferential upregulation of IL-23R in recently divided CLL cells (Fig. 3).

**Fig. 3.**
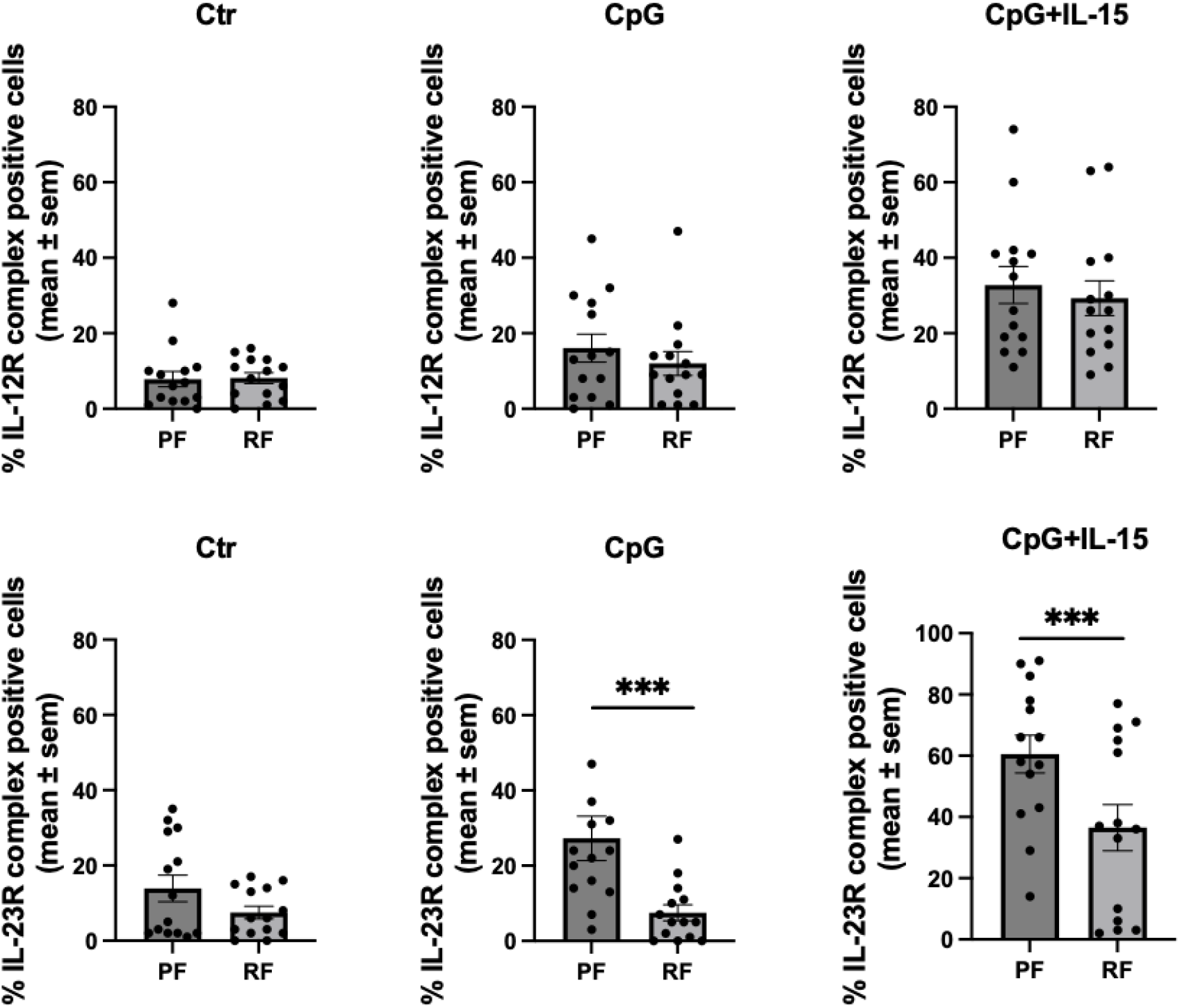
Expression of IL23R and IL12R complex in CLL fractions. A. Expression of IL23R and IL12R complex in PF and RF CLL fractions (n=14) The statistical significance of the difference is evaluated using the two-sided Wilcoxon signed-rank test. *p 0.05; **p 0.01; ***p 0.001; ****p 0.0001

## DISCUSSION

Our study demonstrates that stimulation of CLL cells with CpG and IL-15 results in a selective induction of IL-12 family receptor components. After 72 hours of exposure, a clear upregulation of IL-23R and IL-12Rβ1 was observed, whereas IL-12Rβ2 expression remained largely unchanged. This pattern leads to a robust increase in the expression of the complete IL-23 receptor complex and, to a lesser extent, of the IL-12 receptor complex. The differential regulation of the receptor subunits indicates that innate immune stimulation predominantly primes CLL cells to respond to IL-23 rather than to IL-12, consistent with the notion that IL-23 signaling may be preferentially involved in sustaining inflammatory and survival pathways, while IL-12 responsiveness remains limited due to the weak induction of IL-12Rβ2.

When receptor expression was evaluated in the context of phenotypes that identify CLL intraclonal heterogeneity, defined by CD5 and CXCR4 expression, we observed that IL-23R was more prominently expressed in the proliferative fraction (PF; CXCR4^dim/CD5^bright) compared to the resting fraction (RF; CXCR4^bright/CD5^dim). These findings suggest that IL-23 responsiveness is enriched in the proliferating fraction of the clone, which contains cells recently emigrated from lymphoid proliferation centers. In contrast, the quiescent RF, representing older cells with limited proliferative capacity, showed significantly lower levels of IL-23R. The asymmetric distribution of IL-23R across intraclonal subfractions has several biological implications. First, it reinforces the concept that proliferating CLL cells are susceptible to microenvironmental cytokine signals and may exploit IL-23 signaling to sustain expansion and survival within proliferation centers.

An additional point of relevance is the persistently low expression of IL-12Rβ2 observed in our study. Beyond limiting IL-12 responsiveness in CLL, the absence of IL-12Rβ2 has been mechanistically linked to leukemogenesis. In murine models, genetic deficiency of IL-12Rβ2 predisposes to autoimmunity and spontaneous development of lymphoid malignancies, underscoring a tumor-suppressive role of IL-12 signaling (Airoldi et al., 2005). Thus, the failure of CLL cells to adequately upregulate IL-12Rβ2 in response to CpG + IL-15 stimulation may not only constrain IL-12 signaling but also contribute to creating a permissive context for leukemic persistence and progression. In contrast, the robust induction of IL-23R points to a skewing of cytokine responsiveness toward pro-survival, pro-inflammatory signals, at the expense of the tumor-suppressive functions associated with IL-12Rβ2.

Together, these results refine our understanding of how CLL clones exploit microenvironmental signals. The data support a model in which TLR9- and IL-15–mediated activation enables IL-23 receptor expression on CLL cells, but this occurs in a manner skewed toward the proliferative fraction and coupled with insufficient induction of IL-12Rβ2. From a biological perspective, this asymmetry may contribute to intraclonal specialization, with distinct subfractions of the leukemic population differentially responsive to immune cytokines.

From a therapeutic perspective, our findings suggest two complementary avenues. On one hand, targeting the IL-23/IL-23R axis could be exploited to disrupt pro-survival and proliferative signals, particularly enriched in the proliferative fraction of the clone. On the other hand, strategies aimed at restoring or enhancing IL-12Rβ2 expression and signaling may re-establish the tumor-suppressive functions of IL-12, counteracting the leukemogenic predisposition associated with its absence. The balance between impaired IL-12Rβ2–mediated restraint and enhanced IL-23R–driven activation may therefore represent a central pathogenic mechanism in CLL biology, and future therapeutic interventions could be designed to selectively shift this equilibrium toward an anti-leukemic immune environment.

